# Mitochondria-mitochondria interaction networks show altered topological patterns in Parkinson’s disease

**DOI:** 10.1101/2020.03.09.984195

**Authors:** Massimiliano Zanin, Bruno F. R. Santos, Paul M.A. Antony, Clara Berenguer-Escuder, Simone B. Larsen, Zoé Hanss, Peter Barbuti, Aidos S. Baumuratov, Dajana Grossmann, Christophe Capelle, Joseph Weber, Rudi Balling, Markus Ollert, Rejko Krüger, Nico J. Diederich, Feng Q. He

**Affiliations:** Instituto de Física Interdisciplinar y Sistemas Complejos IFISC (UIB-CSIC), E-07122 Palma de Mallorca, Spain; Center for Biomedical Technology, Universidad Politécnica de Madrid, Campus of Montegancedo, E-28223 Pozuelo de Alarcón, Madrid, Spain; LCSB (Luxembourg Centre for Systems Biomedicine), University of Luxembourg, Campus Belval, 6, Avenue du Swing, L-4367 Belvaux, Luxembourg; Department of Infection and Immunity, Luxembourg Institute of Health (LIH), 29, rue Henri Koch, L-4354 Esch-sur-Alzette, Luxembourg; Centre Hospitalier de Luxembourg (CHL), 4, Rue Nicolas Ernest Barblé, 1210 Luxembourg, Luxembourg; Department of Dermatology and Allergy Center, Odense Research Center for Anaphylaxis (ORCA), University of Southern Denmark, 5000 C, Odense, Denmark; Transversal Translational Medicine, Luxembourg Institute of Health (LIH), 1A-B, rue Thomas Edison, L-1445 Strassen, Luxembourg

**Author notes:** Correspondence should be addressed to F.Q.H.

**Keywords:** Mitochondria-Mitochondria interaction network, network biology, network analysis, scale-free, Parkinson’s disease, neurodegenerative diseases, enteric ganglia, iPSC, Mitochondria

## Abstract

Mitochondrial dysfunction is linked to pathogenesis of Parkinson’s disease (PD). However, individual-mitochondria-based analyses do not show a uniform feature in PD patients. Since mitochondria interact with each other, we hypothesize that PD-related features might exist in topological patterns of mitochondria-mitochondria interaction networks (MINs). Here we showed that MINs form non-classical scale-free supernetworks in colonic ganglia both from healthy controls and PD patients, however, altered topological patterns are observed in PD patients. These patterns highly correlate with PD clinical scores and a machine-learning approach based on the MIN features accurately distinguish between patients and controls with an area-under-curve value of 0.989. The MINs of midbrain dopaminergic neurons (mDANs) derived from several genetic PD patients also display specific changes. CRISPR/CAS9-based genome correction of alpha-synuclein point mutations reverses the changes in MINs of mDANs. Our MIN network analysis opens a new dimension for a deeper characterization of various complex diseases with mitochondrial dysregulation.

## INTRODUCTION

Network-biology approaches are successfully employed for a better understanding of complex diseases that are caused by interactions between genetic and/or environmental factors ^1-6^. Small- and macro-molecules such as genes, proteins and/or metabolites interact with each other and form networks with certain common underlying organization principles in sharp contrast to random networks. All these molecular networks seem to obey to a general scale-free power-law distribution principle^7^, although the definition of power-law distribution might require fine adjustment^8^. Mitochondria, the key organelle regulating cellular metabolism and generating cellular energy, constantly interact with each other, i.e., via the fusion and fission processes. Therefore, they form perpetually changing networks. Nevertheless, it remains unclear whether such organelle interactions form random networks or also well-organized structures obeying to universal principles. Answering this question could open basic new research avenues in neurodegeneration as mitochondrial dysfunction is connected to several neurodegenerative diseases, such as Huntington diseases, Alzheimer’s diseases and Parkinson’s disease (PD) ^9-12^. Therefore, we here took advantage of the availability of various PD-derived tissues and analyzed in all of them whether a functional impairment of mitochondria is associated with any specific topological patterns or features of large-scale mitochondria-mitochondria interaction networks (MINs).

## RESULTS

To obtain more precise information on mitochondria-mitochondria interactions, we used high-resolution 3D mitochondrial immunofluorescence images in colon ascendens (left) and descendens (right) ganglia collected from idiopathic PD patients and healthy controls ^13^. We extracted network adjacency matrixes of mitochondria-mitochondria interactions from all ganglia neurons in such a way that mitochondria were represented as nodes, with an undirected link being present if an interaction is observed between a pair of nodes at the imaging moment. We performed various types of network analyses (up to 19 different network structure metrics) in this work. In a second step, the same network analysis was applied to midbrain dopaminergic neurons (mDANs) differentiated from induced-pluripotent stem cells (iPSCs) derived from skin fibroblasts of genetic PD patients and the corresponding healthy controls. We analyzed the MINs in the samples from patients with heterozygous point mutations, namely in the *alpha-synuclein* (SNCA) gene^14,15^, in the PD-associated gene *RHOT1* encoding a mitochondrial outer membrane GTPase^16,17^ (Miro1), or in the gene encoding the vacuolar protein sorting-associated protein 35 (VPS35)^18,19^. We compared the patients’ samples with that from age- and gender-matched healthy controls as well as mutation-corrected isogenic controls. For the mDANs derived from VPS35-mutated samples, we also analyzed under different culture conditions (with or without anti-oxidants).

### MINs in enteric ganglia neurons form non-classical scale-free supernetworks and are composed of larger subnetworks

As we hypothesized that the universal scale-free principle ^20^ would also apply to mitochondria-mitochondria interaction networks, we first analyzed whether mitochondria form such a network within ganglia. We found that in the MINs, the probability p(k) that a node interacts with k other nodes did not follow a power law ^7^ (i.e., p(k)∼k^−^) (**Fig.1a**). This result indicates that MINs did not self-organize into standard scale-free networks. The inability of MINs to form scale-free networks was independent from the subject groups (PD patients or healthy controls) and from the sample origins (left- or right-side biopsies) (**Fig.1a**). Within the ganglia, the mitochondria formed various sizes of (>16 thousands per group) and types of connected subnetworks/components with different numbers of mitochondria (nodes) and interactions (**Fig.1b**). We therefore checked whether the size of these subnetworks is organized according to a scale-free principle. Interestingly, the overall mitochondria interactions formed a non-classic modular scale-free network, where the probability *P(s)* that one subnetwork with at least *s* nodes indeed decays as a power law, following *P(s)∼k*^*s*^ (**Fig.1c,d**). Unexpectedly, however, the MINs from PD patients were more frequently composed of larger subnetworks than the MINs from healthy controls (*P*-value =10^−17^, see the Methods, **Fig.1c-e**). This difference between healthy controls and PD patients was more evident in the MINs from ganglia out of the right/ascending colon biopsies than that in the left/descending colon (*P*-value =10^−25^, **Fig.1d,e**), possibly due to the assumed rostrocaudal disease progression within the gastrointestinal tract ^21^.

**Figure 1.**
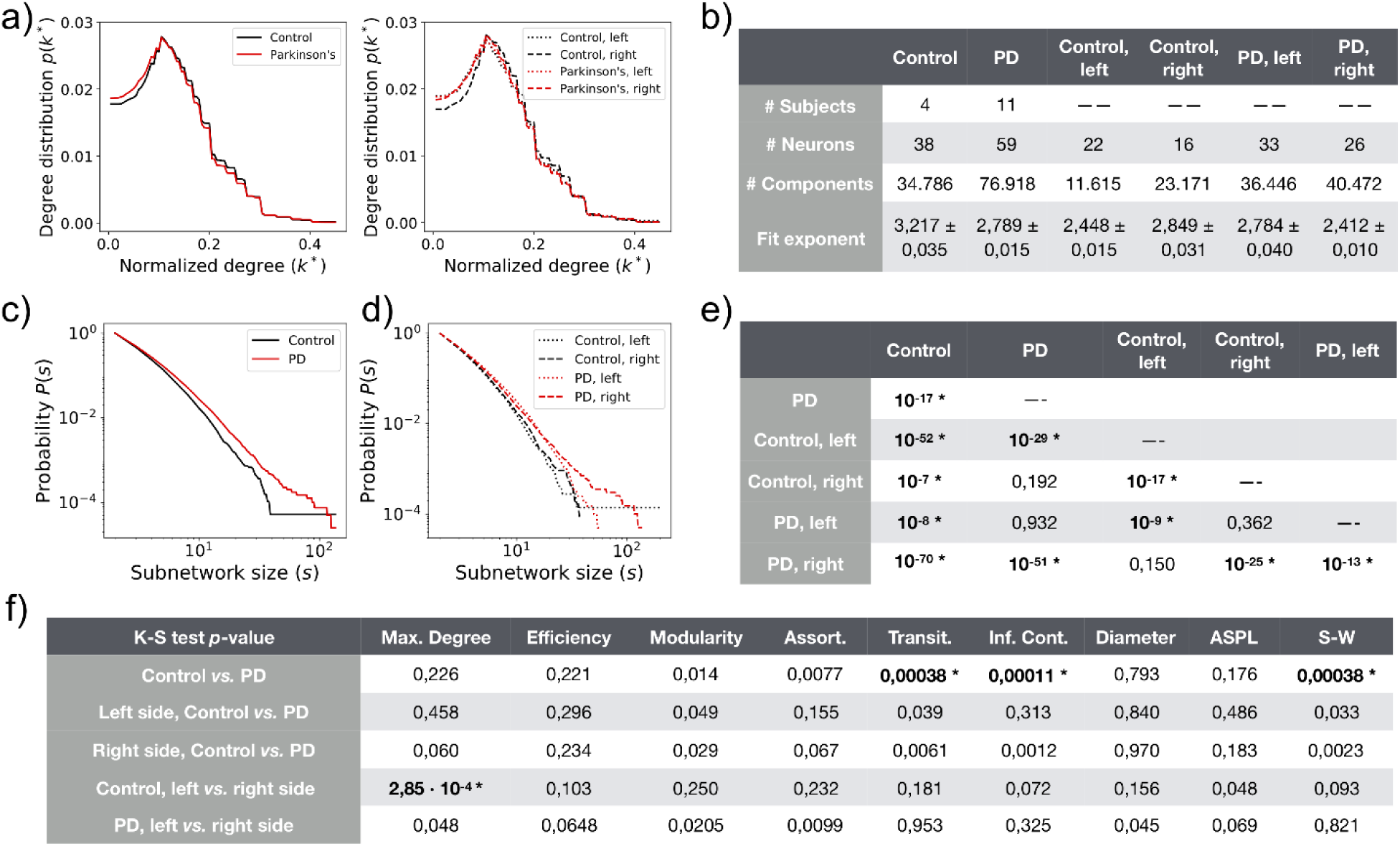
Topological properties of mitochondria interaction networks (MINs) from enteric ganglia of PD. **a**, Probability distribution of the normalized degree of nodes within MINs of PD or healthy controls or indicated subgroups of samples. The displayed degree was normalized by the number of nodes in the given network component. **b**, Summary of the network components of various sets of patients’ materials. A dashed line in the element indicates no entry. **c, d**, Cumulative distribution of the component/subnetwork size of the MINs among all samples of healthy controls and PD patients (c), and among subsets of samples taken from left or right side (**d**). **e, f**, P-value of the test (see the Methods) evaluating the null hypothesis that the exponential fits of the degree distribution for the given two groups share the same power law slope *k* (**e**), or evaluating through a two-sided K-S (Kolmogorov– Smirnov) test the null hypothesis that the distribution of each topological metric of the global MINs is identical for the given two groups (**f**). In **e** only the lower triangular matrix of all the pair-wised comparisons is displayed for simplification. Bold and * indicate a significant P-value<=0.05 after Šidák correction (the displayed P-values are before correction). PD, patients with Parkinson’s Disease; Control, healthy controls; Assort., Assortativity; Transit., Transitivity; Inf.Cont., Information Content; ASPL, average shortest pathway length; S-W, Small-worldness.

### Alteration in network topological features of MINs from enteric ganglia neurons of PD

To systematically explore the topological properties of the MINs, we calculated various other topological/graphic metrics^22^, such as network max degree, diameter, efficiency, average shortest path length (ASPL), modularity, assortativity, transitivity, information content and small worldness (For the definition of different network property metrics, please refer to **Materials and Methods**) ^23^. We first focused on three network metrics with obvious biological meaning and implications. The three parameters including such as network diameter, efficiency and ASPL represent one group of closely related metrics, which all essentially signify how efficiently the energy and information can be transferred and distributed among different nodes/mitochondria within enteric ganglia neurons. The longer the ASPL within a given MIN subnetwork, the less efficiently the MIN subnetwork transfers the energy from one node to another node. Among various analyzed metrics, we only observed marginal differences for transitivity (demonstrating density of triangles), information content (assessing the presence of regular meso-scale structures) ^24^ and small worldness (S-W) between the global MINs from PD patients compared to healthy controls (**Fig.1f and Supplementary Fig.1**). As no topological difference was substantial in the meso-scale properties, we searched for network feature differences at a micro-scale level. Interestingly, for the components with the number of nodes equal to or larger than (≥) 24, we noticed that the average Z-scores of the efficiency were much lower in PD than in healthy controls, whereas that of ASPL were much longer in PD (**Fig. 2a**). In line with this notion, the normalized network diameter for the larger components was much larger in PD than in healthy controls (**Fig. 2a**). These results may explain why PD patients have dysfunctional ganglia neurons ^25^. The observed lower network efficiency and accordingly increased ASPL in MINs might have important implications, i.e., energy and information within enteric ganglia neurons are possibly produced, shared and distributed less competently in the ganglia neurons of PD patients relative to healthy controls. In another hand, similar to that of the power grid ^26^, these network topological features of the MINs may also serve as a protective compensatory mechanism in PD patients. More investigation is required to distinguish between the possible compensatory mechanisms and the pathogenesis causing mechanisms.

**Figure 2.**
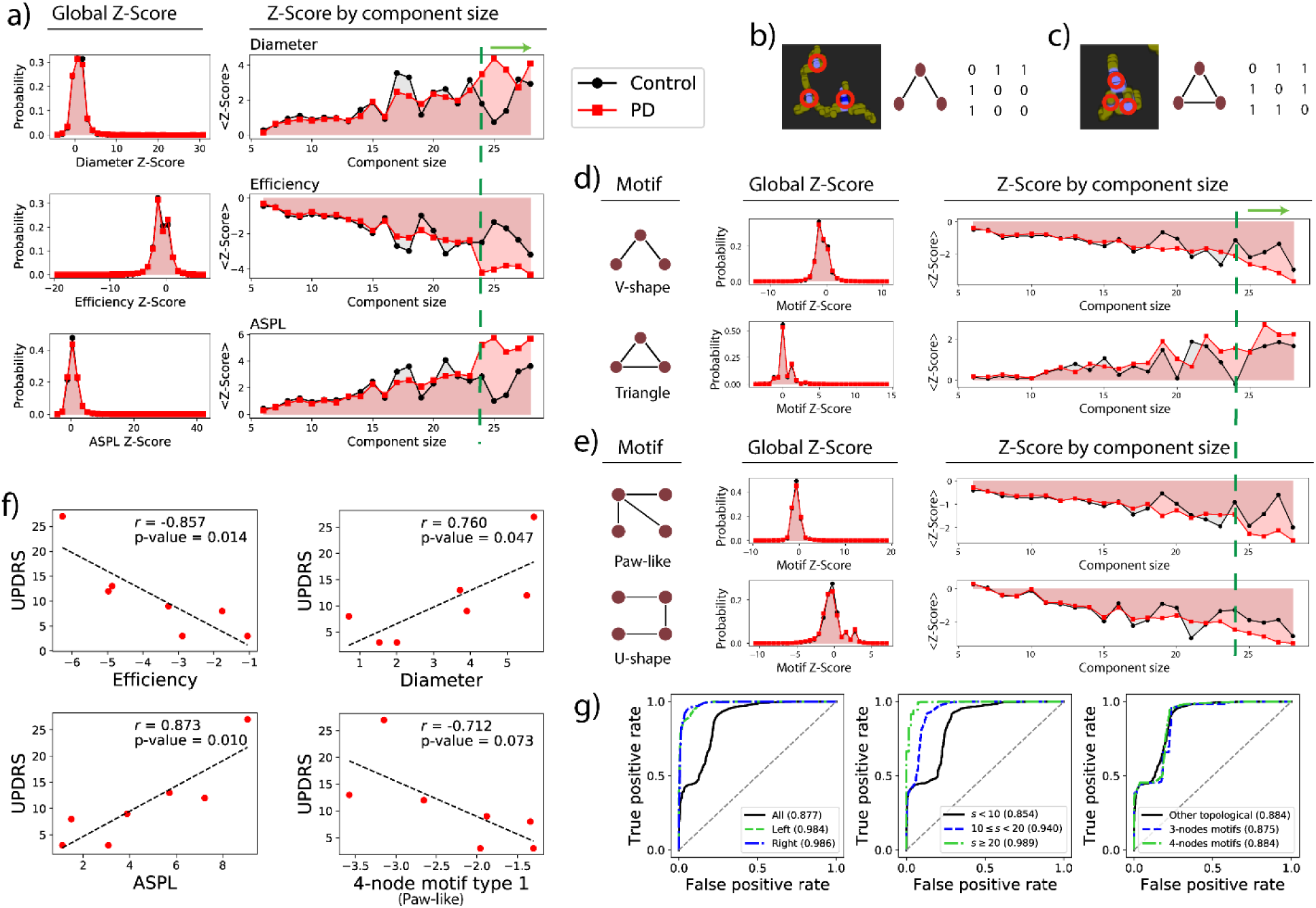
PD MINs show significantly altered motifs and network features and can be used for accurate sample classification. **a**, Distribution of Z-scores of different network metrics. Left, histograms representing the probability distribution of the corresponding Z-score from the global MINs; The right graphs depicting the *Z*-scores for the given size of the MIN components/subnetworks. The green dashed line and the green arrow above the plots highlight the large components. **b, c**, Example of partially- (**b**) or fully- (**c**) connected 3-node MIN motifs. Left, representative mitochondrial immunofluorescence images; middle, the cartoon of the undirected network motif; right, the corresponding adjacency matrix (the pixel values in the link were simplified to 1 if there is a link and otherwise 0). Red circles in the left panel indicates where the nodes are positioned. **d, e**, Distribution of the two types of 3-node (**d**) or 4-node (**e**) motifs from PD patients and healthy controls. Left, the cartoon of the corresponding motif; Middle, the histograms representing the probability distribution of the corresponding Z-scores from healthy controls (black bar) or PD patients (red bar);the right graphs depicting the *Z*-scores for various sizes of the network components. **f**, Correlation analysis between individual patient UPDRS clinical score and network efficiency, diameter, average shortest path length (ASPL) and paw-like 4-node motif (the first one in **e**) within the network components with the size of 28. The parameter r is the Pearson correlation coefficient. P-value is the probability that correlation coefficient is in fact zero. Of note, not all the patients have the corresponding component (size of 28). **g**, Accurate classification of PD patients from healthy controls using the MIN topological features alone. Each graph depicts multiple ROC curves and the corresponding area under ROC curve (AUC). Left, central and right panels respectively present classification results using samples from left- or right-side or both-side ganglia, from components with different sizes, from various indicated features.

Network motifs are recurrent conserved building-blocks composed of a small number of nodes that are often associated with certain functions^27^. Without consideration of the network component size, there was no clear difference in the Z-scores of various types of analyzed motifs between PD patients and healthy controls (for Z-score see Methods, **Fig. 2b-e, Supplementary Fig.2**). Notably, for the components with the number of nodes ≥ 24, we noticed that the partially-connected V-shape 3-node motifs existed less frequently in PD patients than in healthy controls (**Fig. 2d**). This observation seemed to be generally applicable, as it also held true for the partially connected paw-like 4-node motifs (**Fig. 2e**). We also checked other types of MIN motifs and found the fully-connected triangle 3-node motifs possessed much higher Z-score in PD than in healthy controls for large components (size>=24, **Fig. 2d**). This observation was not evident for more-connected 4-node motifs possibly because of the decreased overall frequency of such complex motifs in MINs and randomized networks (**Supplementary Fig.2**). In the mitochondria interacting ‘social’ networks of PD, ‘dysfunctional’ mitochondria relative to ‘normal’ mitochondria might need to fully interact with each other in order to more efficiently maintain the necessary cellular energy supply. Higher frequency of this type of motifs may also partially compensate for the substantial right-side ganglionic shrinking in PD patients ^13^. It is noteworthy that the fully-connected triangle 3-node motifs, as that analyzed in the index of transivity, are the most recurring motifs in many different types of biological and social networks ^28,29^, reflecting the relevance and importance of such types of motifs in establishing network efficiency and maintaining network function.

### Network topological features are correlated with PD clinical scores

With these promising results in terms of difference in topological patterns of the MINs from macro- to meso- to micro-scale levels in mind, we explored whether some of these network features are correlated with the most-used clinical scores, i.e., Unified Parkinson’s Disease Rating Scale (UPDRS) III ^30^ and thus could be used as potential biomarkers of early PD diagnosis. Notably, network efficiency is significantly negatively correlated with the UPDRS scores (r=-0.857, *P-value*=0.014) for a large size of network components (size of 28; this size exists among different individuals), indicating that a lower network efficiency reflects more severe PD motor symptoms in individual patients (**Fig. 2f**). Accordingly, as the related but inverted parameters of network efficiency, ASPL and diameter are highly positively correlated with the UPDRS scores for the corresponding components (size of 28) **(Fig. 2f)**. The Z-scores of the paw-like 4-node network motifs were also negatively correlated with the UPDRS scores, although to a lesser extent **(Fig. 2f)**.

### Network topological features alone can accurately discriminate PD patients from controls

As we had found such a high correlation between network features and well-accepted PD clinical scores, we applied machine-learning approaches to assess whether we can use a combinatory panel of those network features as more powerful biomarkers. After testing and comparing several algorithms in both our real datasets as well as randomized datasets with re-shuffled sample labels **(Supplementary Fig.3**), we selected a high-performance machine learning approach (i.e., multilayer perception (MLP)) with leave-one-out cross-validation to discriminate the samples of PD patients from healthy controls (**Fig. 2g**). When only choosing features from right-side or left-side ganglia, we found the area under the ROC curve (AUC), the essential performance index of biomarkers, was as high as 98.6% and 98.4%, respectively (**Fig. 2g, left panel**). When we mixed the features from both right- and left-side samples, the AUC was still maintained at 87.7% (**Fig. 2g, left panel**). The classification results using various sizes of components of MINs showed that PD-specific features were mainly encoded in large subnetworks (AUC=98.9% for size>=20, **Fig. 2g, middle**). Integration of different types of key topological features is necessary to reach accurate classification (**Fig. 2g**). Together, these results demonstrate that the features in the MINs represent very valuable information and can be used as potent novel biomarkers for the PD diagnosis.

### MINs within dopaminergic neurons derived from genetic PD patients also show altered network features

To further check whether our observation in enteric neurons of idiopathic PD patients holds true in mDANs derived from genetic PD patients, we generated human iPSCs-differentiated mDANs and analyzed their MINs (**Fig. 3a**). Again, the MINs in iPSCs-differentiated mDANs, no matter being derived from which genetic PD patients or age- and gender-matched healthy controls, did not self-organize into standard scale-free networks similar as observed in enteric ganglia neurons (**Fig. 3b**). In line with the observation in enteric ganglia, the MINs of iPSCs-differentiated mDANs derived from a PD patient with a point mutation in the *SNCA* gene that leads to an A30P amino acid exchange in the encoded protein also formed a non-classic scale-free supernetwork (**Fig. 3c**). Consistent with the effect on subnetwork sizes of the MINs from the idiopathic PD patients, the SNCA-mutated patient showed much larger subnetworks than that from the healthy controls (50 or 60 different measurements or clones per group; *P*-value=3.81×10^−13^, **Fig. 3c**). This also holds true for that from one Miro1-mutated patient relative to the age- and gender-matched healthy control (*P-value*=7.06 ×10^−44^). However, the correction of the point mutation in SNCA using CRISPR/CAS9-based genome editing did not dramatically, although still significantly (*P-value*=1.65 ×10^−3^), change the size of the subnetworks (**Fig. 3c**). In contrast to the observations of other mutations, the VPS35-mutated patient showed substantially smaller subnetworks than that from the matched healthy controls (*P-value*=2.15×10^−53^, **Fig. 3c**), which is in line with the reported observation that VPS35 mutations cause the fragmentation of individual mitochondria^19^. To further test whether the astonishing effect of the VPS35 mutations is regulated by oxidative stress, we exposed the cells to oxidative stress during the iPSCs-differentiation process. Following oxidative stress, the difference disappeared in subnetwork size of MINs within mDANs derived from the VPS35-mutated patient versus (vs.) the matched control, indicative of the direct involvement of oxidative stress in VPS35-mediated feature changes of the MINs. The distinction observed between the VPS35-mutated samples and the other genetic PD samples might be simply attributable to the fact that the D620N mutation in VPS35 disrupts both the distribution of endosomes^31^ and mitochondrial functions^19^, while the other PD genetic mutations mainly affect the latter. Therefore, the effect on subnetwork size of MINs of mDANs might be dependent on which familial PD gene is mutated, depending on whether the given mutation directly contributes to mitochondrial dysfunction and is regulated by oxidative stress.

**Figure 3.**
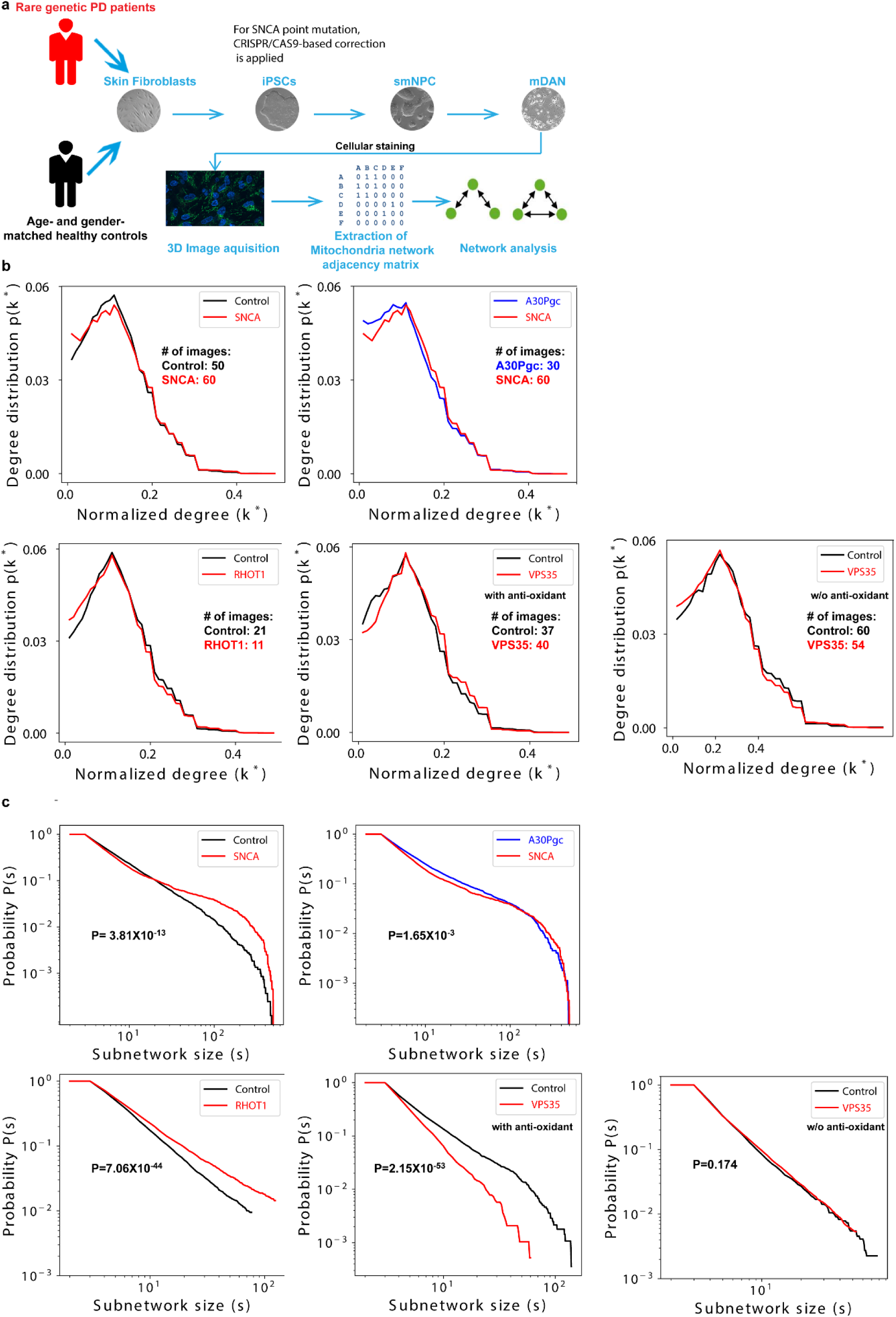
MINs within iPSCs-differentiated dopaminergic neurons of certain genetic PD patients show similar alterations as that in ganglia of sporadic PD. **a**, Schematic on how to perform network analysis from dopaminergic neurons differentiated and derived from skin fibroblasts of human subjects. Details are provided in the Materials and Methods. Here we only described briefly. iPSCs were first reprogrammed from skin fibroblasts using Yamanaka factors. Using small molecules, we then differentiated iPSCs to small molecule neural precursor cells (smNPCs). Finally, using trophic factors, smNPCs were differentiated to midbrain dopaminergic neurons (mDANs). We then performed cellular staining to identify mitochondria within mDAN and identified mitochondria-mitochondria interactions using confocal 3D mitochondrial immunofluorescence images. After extracting network adjacency matrixes, we then performed large-scale network analysis on the MINs. **b**, Probability distribution of the normalized degree of nodes within MINs of iPSCs-differentiated mDANs derived from skin fibroblasts of different genetic PD patients or the corresponding age- and gender-matched healthy controls (refer to Materials and Methods) or SNCA-mutations corrected lines using the CRISPR/CAS9 approach. Of note, mDANs from the VPS35-mutated patient and the matched controls were differentiated with or without (w/o) anti-oxidants (as indicated), while the others were all differentiated in the presence of anti-oxidants. **c**, Cumulative distribution of the component/subnetwork size of the MINs among all the samples of different patients or controls or patients’ isogenic controls. Displayed *P*-value of the test (see the Methods) evaluating the null hypothesis that the exponential fits of the degree distribution for the indicated two groups of the given plot share the same power law slope *k.*

To obtain a more comprehensive understanding of the network features of MINs, we investigated and compared other network topological indexes of the MINs from mDANs derived from genetic PD patients. Keeping in mind the observations in ganglia neurons, we particularly checked the topological metrics related to network efficiency. Notably, the MINs from the three rare genetic PD patients all presented smaller diameters for the subnetworks that are composed of nodes larger than a certain number (≥34, 27, 34, 14 for SNCA mutation, RHOT1 mutation, VPS35 mutation with or without oxidative stress respectively, **Fig. 4a**), whereas the efficiency was always higher than that of age- and gender-matched healthy controls (**Fig. 4b**). Correction in the SNCA mutation reversed both changes in network diameters and efficiency caused by the SNCA mutation (**Fig. 4a, b**). As determined by the definition of ASPL, the change of the ASPL in genetic PD patients is correlated with that of network diameter (**Supplementary Fig. 4a**). Of note, again the effect of the VPS35 mutation on these MIN indexes, when the differentiation was performed with anti-oxidants, was smaller compared with that of the other analyzed genetic factors in this work. As oxidative stress worsens the iPSC-derived mDAN phenotypes of several genetic factors that contribute to the pathogenesis of PD^32,33^, the influence of the VPS35 mutation under oxidative stress on particular network indexes became more evident (**Fig. 4a, b and Supplementary Fig. 4a**). In short, closely associated network indexes analyzed here such as diameter, efficiency and ASPL showed consistent alteration in all the selected genetic PD patients. The results of the three network parameters demonstrated that the MINs within mDANs derived from several genetic PD patients all have enhanced efficiency in terms of energy transfer among different mitochondria within the those larger subnetworks. Most likely, these consistent alterations in particular network features represent a conservative compensatory mechanism that tends to protect PD mDANs from death.

**Figure 4.**
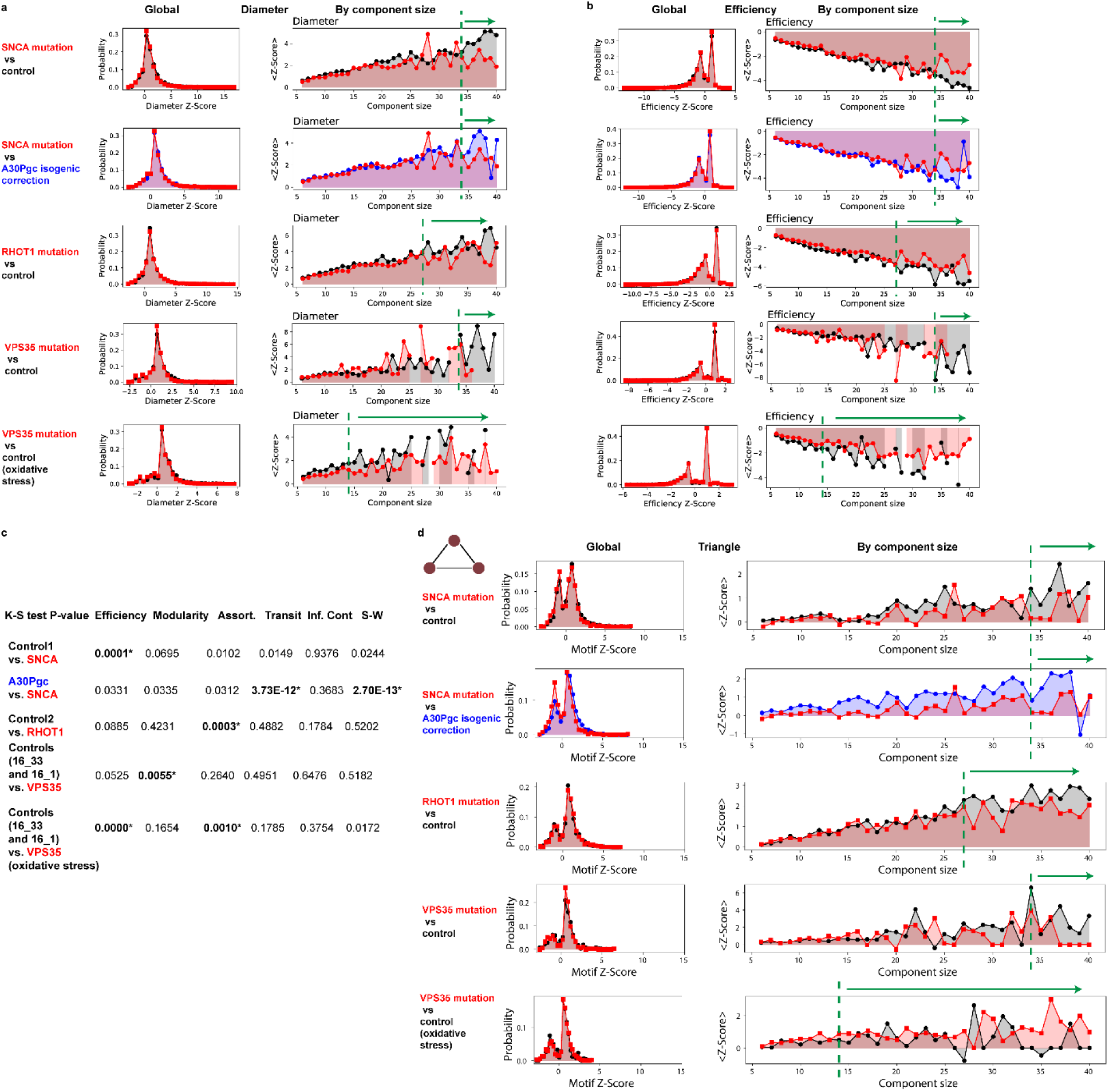
MINs of iPSCs-differentiated mDANs of different genetic PD patients show consistent alteration in network features. **a, b**, Distribution of Z-scores of different network structure features, for instance network diameter (**a**) and efficiency (**b**). Left, histograms representing the probability distribution of the corresponding Z-score from the global MINs of mDANs derived from different patients or matched controls; The right graphs depicting the *Z*-scores for the given size of the MIN components/subnetworks. The green dashed line and the green arrow above the plots highlight the large components that show a clear difference. **c**, *P*-values of the two-sample two-sided K-S (Kolmogorov–Smirnov) test evaluating the null hypothesis that the distribution of each indicated topological metric of the global MINs is identical for the given two groups. Bold and * indicate a significant P-value<=0.05 after Šidák correction (the displayed P-values are before correction). **d**, Distribution of the fully-connected 3-node MIN motifs (as known as ‘triangle’) of mDANs derived from different patients or matched controls; Left, the histograms representing the probability distribution of the corresponding Z-scores of global MINs from different genetic PD patients or matched controls; the right graphs depicting the *Z*-scores for various sizes of the network components.

We further analyzed other topological indexes of MINs of mDANs that were also calculated in enteric ganglia neurons of idiopathic PD patients. Interestingly, correction in the SNCA mutation significantly affected network transitivity and small worldness (**Fig. 4c**, for transitivity and small worldness, refer to the **Materials and Methods** for the definition). The MINs of mDANs derived from both VPS35-mutated materials under oxidative stress and Miro1-mutated patient samples showed significantly changed assortativity, a network metric representing to which extent highly connected nodes in a network tend to link with each other^34^ (**Fig. 4c**, refer to Methods). Furthermore, the MINs of mDANs derived from both VPS35-mutated materials under oxidative stress and SNCA-mutated materials showed a significant change in the MIN network efficiency even in a global scale (**Fig. 4c**) in addition to those visible only in larger subnetworks (**Fig. 4b**). For the mDANs derived from the VPS35-mutated patient materials cultured with anti-oxidants, only modularity of the global MINs that measures how much the network is organized into communities showed a significant difference (**Fig. 4c**). Thus, alike the effect on MIN subnetwork size, the influence on particular network properties is also dependent on specific genetic mutations and apparently affected by exogenous oxidative stress.

We further examined the 3-node and 4-node motifs of MINs from those genetic PD patients. Interestingly, the triangle motifs, the most abundant network motifs in different types of networks^28,29^, within the mDANs differentiated with anti-oxidants also displayed similar changes for larger MIN subnetworks of all patients carrying SNCA- or RHOT1- or VPS35-mutations (**Fig. 4d**). Since both cellular MINs and power/electricity grid networks^29^ might share similar functions in terms of ‘energy transferring’, we reasoned that the frequency reduction in the triangle motifs of the larger MIN subnetworks from those genetic PD patients might enhance the risk of energy supply failure and eventually harm the functions and survival of those neurons. The correction in the SNCA point mutation again reversed the frequency change of triangle motifs caused by the SNCA mutation (**Fig. 4d**). It is noteworthy that under oxidative stress the MINs derived from the VPS35-mutated patient showed an inverted change as that of mDANs derived from the PD patients with mutations in any of the three analyzed key PD genes, when being differentiated in the presence of anti-oxidants. This oxidative-stress induced effect of the VPS35 mutation on the frequency of triangle motifs of iPSCs-differentiated mDANs was in fact similar to that observed in the *ex-vivo* analysis of ganglia neurons of idiopathic PD patients (**Fig. 2d**). These results are very much in line with the current well-established paradigm that oxidative stress plays a critical role in dopaminergic neurotoxicity^35^ (**Fig. 4d**). The frequency change of the V-shape 3-node motifs was similar, although to a lesser degree, as that of triangle motifs (**Supplementary Fig. 4b**). The frequency change of both paw-like and U-shape 4-node motifs in those genetic PD patients was still similar to that of the triangle 3-node motifs in larger MIN subnetworks (**Supplementary Fig. 4c, d**). Nevertheless, the altered degree between the genetic PD patients and the controls in the frequency of the analyzed 4-node motifs was smaller compared with that of the triangle motifs. In summary, the analyzed network motifs also showed consistent changes for four out of the five comparison groups/conditions. The only exception existed in the MIN network features caused by the VPS35 mutations that was imposed by oxidative stress. Taken together, although not always identical in changes for various examined indexes, the image datasets from both idiopathic and genetic PD patients revealed novel critical changes in the topological structure of MINs that are associated with PD.

## DISCUSSION

Since mitochondria constantly interact with each other, it is rational to analyze the mitochondria-mitochondria interaction networks (MINs). However, such an analysis has never been performed even in general populations, not to mention among the patients with neurodegenerative diseases with direct mitochondrial involvement. As many molecular networks share certain underlying organizing principles, we aimed to investigate whether the MINs follow similar principles and whether and how the network topological properties are affected in relevant pathological conditions that are related to mitochondrial deficiency. To this end, we here comprehensively analyzed a variety of network topological indexes of MINs, contrary to a conventional analysis focusing on individual-mitochondria-based phenotypes such as mitochondrial number, volume, size, shape and even recent simplified network-like analysis only on connection degree ^13,36^. Beyond the initially only intended proof-of-principle analysis, we found remarkable pattern differences in the MINs of enteric ganglia of sporadic PD patients vs. healthy controls. Furthermore, particular network metrics are highly correlated with PD clinical scores, indicating a potential of using particular network features for early diagnosis and basic research purposes of PD. Excitingly, with network topological features alone, we can already accurately distinguish the PD patients from healthy controls. This discovery opens the door to a new type of biomarkers from network metrics of MINs in patient-based materials. However further validation in a large-scale cohort or even multicenter cohorts is required. In PD patients these differences in MINs might be directly related to well-known mitochondrial complex I deficiency ^37^, mitochondrial fragmentation and/or deficient mitochondrial dynamics ^38,39^. In this work, we demonstrated the association between altered network topological indexes of MINs with known mitochondrial deficiency of sporadic PD patients.

Network analysis of MINs of mDANs derived from all the tested genetic PD patients vs. age- and gender-matched healthy controls revealed consistent changes for several related network metrics, such as diameter, efficiency and ASPL. Remarkably, the change of direction seen in genetic PD patients is in sharp contrast to that seen in sporadic PD patients. The difference in change direction of particular network features might be caused by several factors: 1, genetic PD vs. idiopathic PD; 2, tissue difference (enteric neurons vs. mDANs); 3, ex vivo imaging in ganglia vs. imaging on in vitro derived cells; 4, also possibly direct disease involvement vs. secondary compensation mechanism. Therefore, the inverted change of direction is plausible due to these fundamental differences. Importantly, despite of a huge difference in the roles of the tested genetic factors, the consistence in pattern changes of particular network indexes (e.g., network efficiency related indexes and triangle motifs) among different genetic PD patients underscores the value of using this type of MIN network analysis to assist diagnosis and classification of genetic PD patients. Such a consistency in pathology among different genetic PD patients has so far not been reported in other studies not applying such a fundamental network analysis in MINs. Machine-learning-based computational analysis of MINs provides another layer of new information and enables automatized classification of a large number of subjects.

We also noticed a general negative correlation between the changed directions (increase or decrease) in network efficiency and triangle motif frequency of PD patients, independent of samples from sporadic or genetic PD patients. These two important network metrics might well compensate each other to fine tune the overall functions of mitochondria networks in PD patients no matter in which tissue we analyzed. Interestingly, the only exception existed in the MINs of the mDANs derived from the VPS35-mutated patient materials under oxidative stress. In that case, both network efficiency and triangle motif frequency in the MINs were simultaneously heightened, possibly to fight against or compensate the cytotoxic effects induced by oxidative stress.

Due to the limited access to colonic ganglia samples from healthy controls, we were unable to analyze more healthy controls at the current stage of the project. We also could not access the ganglia materials from patients with other types of diseases, in particular from other (e.g., inflammatory) colon diseases. Due to this lack of comparison with samples from non-PD patients, we cannot conclude whether our observed changes in the structural features of the MINs are PD-specific or not. However, we are poised that such MIN-related network analyses provide novel insight into the pathogenesis and/or compensatory mechanisms in various chronic complex diseases. The MIN signature, *per se*, could be qualified as a key health indicator, providing information on the energy supply (deficits) in various diseases. Such analyses open innovative avenues of biomedical research for dissecting complex diseases, with primary or secondary bioenergetic deficiencies. Finally, this approach may well be applicable to the network analysis of other cellular organelles, such as endoplasmic reticula or lysosomes.

## MATERIALS AND METHODS

### Experimental Methods

#### Reprogramming of human fibroblasts into iPSCs

We complied with all the relevant ethic regulations and Luxembourg CNER (Comité National d’Ethique de Recherche) has approved the usage of the iPSCs derived from PD patients and the related controls (201411/05). Both the SNCA-mutated patient (p.A30P) and the unaffected control (Control 1) were 67-year old male when the biopsies were collected. The RHOT1-mutated patient was from the existing German PD cohort (average age of onset of 59.4 ± 13.2 years, average age of sample collection of 65.7 ± 10.2 years). Informed consent was obtained from these patients and controls and approved by the Ethics Committee of the Medical faculty and the University Hospital Tübingen, Germany. The RHOT1-mutated late-onset female PD patient (with a heterozygous point mutation c.815G>A in RHOT1 [NM_001033568]) had a tremor dominant clinical phenotype and her father had also tremor in family history. The selected control (Control 2) was age- and gender-matched. More information about the RHOT1-mutated patient can be found elsewhere^17^.

Patient dermal fibroblasts carrying the heterozygous p.D620N mutation in VPS35 were a kind gift from George Mellick from the Griffith Institute (Queensland, Australia). Control fibroblasts from age and gender-matched healthy individuals 16_33 and 16_1 are from Tübingen’s Biobank. Skin biopsies were performed at the ages of 73, 72 and 77 for VPS35-mutated patient, the control 16_33 and the control 16_1 respectively. Informed consent were given by all individuals included in this study.

Skin fibroblasts of patients or healthy controls were cultured at low passage number and maintained with Dulbecco’s modified eagle medium (41965-062, Thermo Fisher Scientific) supplemented with 15% fetal bovine serum (10270106, Thermo Fisher Scientific) and 1% penicillin-streptomycin (15140-122, Thermo Fisher Scientific). When confluency was reached, wild-type skin fibroblasts were reprogrammed into induced-pluripotent stem cells (iPSCs) via lentivirus infection ^40^ using the CytoTune-iPS 2.0 Sendai Reprogramming Kit (A16517, Thermo Fisher Scientific) and patient-derived fibroblasts were reprogrammed into iPSCs via synthetic modified mRNA ^41^. For the samples derived from control 16_1, the fibroblasts were reprogrammed using the three plasmids (pCXLE-hOct3/4 [Addgene #27076], pCXLE-hSK [Addgene #27078], pCXLE-hUL [Addgene #27080]) with 10ug of each plasmid through the Amaxa Nucleofector (Lonza). The fibroblasts from other donors were reprogrammed using Sendai virus.

iPSC clones were expanded in culture and maintained with Essential 8 medium (A1517001, Thermo Fisher Scientific) supplemented with 1% penicillin-streptomycin. Chosen iPSC clones for neuronal differentiation were selected via karyotype analysis and iPSC-characterization procedures^42^.

#### Midbrain dopaminergic neuronal differentiation

Following the procedures above, human iPSCs derived from patients or age- and gender-matched healthy controls were obtained and submitted to neuronal differentiation (for details see below). We included human iPSCs from a monogenic, heterozygous dominant familial case of PD, with a point mutation in the *alpha-synuclein* (SNCA) gene (Patient 1), from a healthy control (Control 1), and from a patient isogenic control (Patient1 + mutation correction). The patient isogenic control was obtained by the CRISPR/CAS9-based genome editing to correct the point mutation of SNCA found in the Patient 1 case. The detailed method was described elsewhere ^43^. We also generated human iPSCs from a genetic PD patient with a point mutation in RHOT1 (Patient 2), from a matched healthy control (Control 2), from a familial genetic PD patient with a point mutation in VPS35 and from two matched healthy controls (Control 16_33 and 16_1).

Chosen iPSC clones were differentiated into small-molecule neural progenitor cells (smNPCs) via small molecules of human neural progenitors ^44^. Successfully differentiated smNPCs were expanded in culture and maintained with N2B27 medium consisted of 50:50 Neurobasal (21103-049, Thermo Fisher Scientific)/DMEM-F12 (11320-033, Thermo Fisher Scientific) supplemented with 1:200 N2 (17502-048, Thermo Fisher Scientific), 1:100 B27 (17504-044, Thermo Fisher Scientific), 1% Glutamax (35050-061, Thermo Fisher Scientific) and 1% penicillin-streptomycin. Dopaminergic neuronal differentiation of smNPCs was performed using the methodology explained elsewhere ^44^. Of note, the cells derived from the VPS35-mutated patient and the matched controls were differentiated in the medium with or without N2 supplements (as anti-oxidants), while all the others were cultured with N2 supplements.

#### Live-cell imaging of iPSC-derived neurons

For the materials derived from the patient with a mutation in RHOT1 or the related control, neurons at day 25 of maturation were seeded in chamber slides (154534, Thermo Fisher Scientific). Five days were needed for the cells to stabilize in the chamber slides and reach an appropriate level of connectivity. For the materials derived from the patient with a mutation in VPS35 or the related control, neurons at day 21 of maturation were seeded in chamber slides (154534, Thermo Fisher Scientific). Nine days were needed for the cells to stabilize in the chamber slides and reach an appropriate level of connectivity. For the cells derived from both the RHOT1-mutated patient and the VPS35-mutated patient and their corresponding controls, the staining and image analysis is identical. At day 30 of maturation, mDANs were stained for live-cell imaging by using 1:10000 MitoTracker Green FM (M-7514, Thermo Fisher Scientific) to label mitochondria and 1:5000 LysoTracker Deep Red (L-12492, Thermo Fisher Scientific) to label lysosomes. Cellular nuclei were stained with 1:100 Hoechst 33342 (H1399, Thermo Fisher Scientific) after mitochondria and lysosomes staining was performed. Neurons were washed once with pre-warmed medium prior to imaging. Live-cell imaging was performed using the Live Cell Microscope Axiovert 2000 with spinning disc (Carl Zeiss Microimaging GmbH) in a humidified atmosphere containing 5% CO2 at 37°C.

For the cells derived from the alpha-synuclein (*SNCA*)-mutated patient or the corresponding control, the details were slightly different. Neurons at day 35 of differentiation were seeded into coverslips (AB0577, Thermo Fisher Scientif). Ten days were necessary for the cells to stabilize and regenerate the complex network. At day 45 of differentiation, cells were fixed using 4% PFA (J61899.AP, Thermo Fisher Scientific) for 15 minutes at room temperature agitating. Permeabilisation/blocking was performed using 0.4% Triton-X-100 (T8787-100ML, Sigma-Aldrich) in PBS +/+ (HYCLSH30264.FS, GE Healthcare Europe GmbH) with 10% Goat serum (S26-100ML, Merck Millipore) and 2% BSA (B9000S, New England Biolabs) for 1 hour at room temperature. First and second antibodies were prepared in a solution of 0.1% Triton-X-100 (T8787-100ML, Sigma-Aldrich) in PBS+/+ (HYCLSH30264.FS, GE Healthcare Europe GmbH) with 1% Goat serum (S26-100ML, Merck Millipore) and 0.2% BSA (B9000S, New England Biolabs). For mitochondria detection, we used the Tom20 (sc-11415, Santa Cruz) antibody at 1:500 dilution overnight at 4°C with agitation. Tom20 was detected by the use of the secondary ab Alexa Fluor® 488 (A-11008, Thermo Fisher Scientific) at 1:1000 dilution, incubated for 3 hours at room temperature with agitation. For nuclear staining we used 1:100 Hoechst 33342 (H1399, Thermo Fisher Scientific) for 15 minutes at room temperature with agitation. Coverslips were mounted in slides using Vectashield (H-1000, LABCONSULT SPRL / Vectorlab) mounting solution and sealed. Imaging was performed using the Live Cell Microscope Axiovert 2000 with spinning disc (Carl Zeiss Microimaging GmbH) using a 63x objective. For each condition, it was acquired ten non-empty fields randomly selected, each of them as a Z-stack, using a 0.2 µm Z-axis step and a total number of slices enough to cover the entire depth of the sample. All files were saved for further analysis as .czi files.

### Computational Methods

#### MIN matrix construction

Adjacency matrices of the mitochondrial interaction networks (MINs) were extracted from confocal three dimensional (3D, with Z-stack) mitochondrial immunofluorescence images of colonic ganglia ^13^, according to a reported method which has been optimized for image-based network analysis^45^. In classical adjacency matrices (A) of undirected graphs, the element A_i,j_=1 indicates that there is a link between nodes i and j, A_i,j_=0 otherwise. In contrast to the classical matrix, in the adjacency matrix variant defined by Kerschnitzki et al. ^45^, the matrix element A_i,j_ represents the count of pixels in the link connecting the given nodes i and j if there is a link and otherwise sets to zero. The key Matlab functions for mask skeletonization and adjacency matrix extraction, namely ‘Skeleton3D’ and ‘Skel2Graph3D’ were kindly provided by the authors of the previous work^45^. For the parameter ‘THR’ of the function ‘Skel2Graph3D’, defining the minimum length of branches (edges that do not end at another node), to filter out potential skeletonization artifacts, in our analysis we set as zero to avoid losing any information. Accordingly, in the following network analysis, we considered the existence of a link between the nodes i and j if A_i,j_ is larger than zero. The other criteria, masks and filters used for mitochondrial segmentation and pixel calculation were described in our previous work ^13^.

#### Subnetwork/component extraction and computational analysis

The MINs, reconstructed as aforementioned, are potentially disconnected, i.e., they may not form a path between all pairs of nodes. In order to ensure a meaningful calculation of all the analyzed topological metrics, we have proceeded to dissect each MIN into a collection of connected subnetworks/components, thus representing a set of locally interacting mitochondria.

The following standard metrics have been evaluated on the obtained subnetworks:

- ***Normalized degree (k*).*** Considering the varying sizes of distinct network components and to make the degree comparable, we normalized the connection degrees of the given nodes within each component using the equation 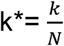, where *k* is the raw connection degree and *N* is the number of nodes in the given component/subnetwork.
- ***Link density and Max Degree.*** Respectively defined as the number of active links over the total number of possible links 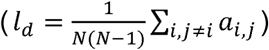, and the number of direct connections of the most connected node (*m*_*d*_ *= max*_*i*_ *k*_*i*_) ^7,22^.
- ***ASPL.*** The average shortest path length (ASPL) is defined as the average distance separating all possible pairs of nodes, i.e., 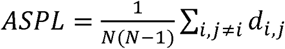^22^.
- ***Diameter.*** Defined as the greatest distance between all pairs of nodes in the network^22^.
- ***Efficiency***. Metric assessing how efficiently information can be transmitted among nodes; it is defined as the harmonic mean of the geodesic distance between all pairs of nodes: 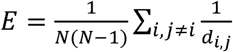, *N* being the number of nodes composing the network, and *d*_*i,j*_ the distance (in terms of the number of links) between nodes *i* and *j* ^46^.
- ***Modularity***. Measuring how much the network is organized into communities, i.e., groups of nodes strongly connected between them but loosely connected with the other nodes of the network ^47,48^. The community structure has been detected through the Louvain algorithm ^49^.
- ***Assortativity***. Pearson’s correlation coefficient between the degrees of both nodes of a link. Positive values indicate that highly connected nodes prefer to link with other hubs while negative values designate that highly connected nodes prefer to link with periphery nodes^34^.
- ***Transitivity***. Density of triangles (triplets of completely connected nodes) in the network.
- ***Information Content***. The measure of assessing the presence of meso-scale structures, e.g. communities, based on the identification of regular patterns in the adjacency matrix of the network, and on the calculation of the quantity of information lost when pairs of nodes are iteratively merged ^24^.
- ***Small-Worldness***. Metric assessing the coexistence of a high clustering coefficient and a low mean geodesic distance ^23,50-52^.
- ***Motifs***. Specific connectivity patterns, created by a small number of nodes, that exist more frequently in the given networks than in randomized networks ^27^. Motifs with 3 or 4 nodes have been considered here. We displayed the components with a size ranging from 6-28 **Fig2d, e**. The component with 29 nodes only appeared once in the healthy controls. When showing the average scores, we did not include very large components (size>=29) in **Fig2d, e**. This restriction in sizes does not apply to the classification section of ganglia samples.
- ***Comparison of subnetwork size between groups.*** Distribution of network components’ sizes, e.g. the curves in **Fig.1b**, have further been modelled through a power law *P(s)∼k*^*s*^, where *s* is the component size, by disregarding the lower and higher tails of the curves ^53^. Pairs of distributions have been compared for the null hypothesis of sharing the same power law slope *k* (**Fig. 1e and 3c**), by representing the two slopes as two random variables from two normal distributions, and by calculating the probability of the difference between the means of both distributions of being zero.

#### Normalization through random networks

In order to normalize the values obtained for the listed metrics, a set of 100 random networks were generated for each component, and used as a null model. Each one of these randomized networks is composed of the same number of nodes and links as the original network; additionally, to ensure a biological plausibility, each generated random network was used only if all the nodes and links form a single component. Afterwards, each metric is normalized through a Z-score, calculated as: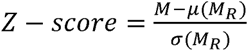, *M* being the value obtained in the real network, *M*_*R*_ the set of values obtained for the random set, and *µ*(*·*) and *σ*(*·*) respectively the average and standard deviation operators.

##### Probability of overall Z-scores

For any of the analyzed metrics, the probability of a given Z-score is defined as follows: 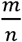, where *m* is number of components with the given Z-score and *n* is number of total components within the MINs.

#### Classification

The classification models’ performance has been corrected against overfitting by using a modified Leave-One-Out Cross Validation (LOOCV) approach. The standard LOOCV technique entails testing each instance of the data set with a model trained with all other instances, followed by calculating the average classification error. It is worth noting that a simple LOOCV would here lead to an overfitting, as each person in the data set is described by multiple instances (i.e. different neurons and mitochondrial networks). False conclusions may be drawn when using a model trained from the MINs of one neuron for testing another neuron of the same participant. In order to avoid this pitfall, we here employed a modified approach in which each model was trained using the data from all the other people, except from those records belonging to the tested participant. The overfitting issue was also minimized by the fact that we had many more network components/subnetworks than the selected features. Furthermore, we also randomly re-shuffled the sample labels 50 times to test whether the high AUC values we achieved in the real datasets can be also obtained even in the randomized datasets.

### Quantification and Statistical Analysis

We employed the two-sample two-sided K-S (Kolmogorov–Smirnov) test in general. However, for the comparison of the exponential fits (**Fig. 1e and 3c**), we used a different test as a K-S test would not work well with a distribution with such a long tail (for details see the description above). Bold and * indicate a significant P-value (<=0.05) after Šidák correction (the displayed P-values in the corresponding figures are before correction). Whenever the corresponding test was used, it was directly indicated in the corresponding figure legend. The number of analyzed samples were indicated either in Fig.1 or in Fig. 3.

### Data and Code Availability

Raw 3-D image datasets will be deposited online (>500G; it will take time to deposit everything) and the computational script codes can be accessible at *Githu*b (https://github.com/FengHe001/Mitochondria-network-analysis; https://github.com/FengHe001/Network-matrix-extraction). Network adjacency matrix was extracted using *Matlab* codes and network analysis was performed using *Python* scripts. All these information will become openly accessible upon acceptance.

## ACKNOWLEDGEMENTS

F.H. was partially supported by Luxembourg National Research Fund (FNR) CORE programme grant (CORE/14/BM/8231540/GeDES), FNR AFR-RIKEN bilateral programme (TregBAR, F.H. and M.O.), PRIDE programme grants (PRIDE/11012546/NEXTIMMUNE and PRIDE/10907093/CRITICS). The work was also partially supported through intramural funding of LIH and LCSB through Ministry of Higher Education and Research (MESR) of Luxembourg. The cooperation was achieved through the European Cooperation in Science and Technology (eCOST) Action CA15120 OpenMultiMed. Fibroblasts were obtained from the Neuro-Biobank of the University of Tübingen, Germany. This biobank is supported by the local University, the Hertie Institute and the DZNE.

## AUTHOR CONTRIBUTIONS

M.Z. performed all the computational analyses and wrote the manuscript. B.F.R.D.S., P.B., C.B.E., S.B.L. D.G., Z.H., performed iPSCs-related experiments. P.A. constructed the adjacency matrix. C.C. performed parts of analyses. J.W. obtained the biopsies. P.A. and A.B. prepared and provided the imaging materials of ganglia. R.B., M.O., R.K, and N.D. provided substantial insights and supervision into the project. F.H. conceived and oversaw the whole project and wrote and revised the manuscript.

## DECLARATION OF INTERESTS

The authors declare no competing interests.

## SUPPLEMENTARY INFORMATION

Supplementary figures are directly attached in the end for the initial submission.

## SUPPLEMENTARY FIGURES

**Supplementary Figure 1.**
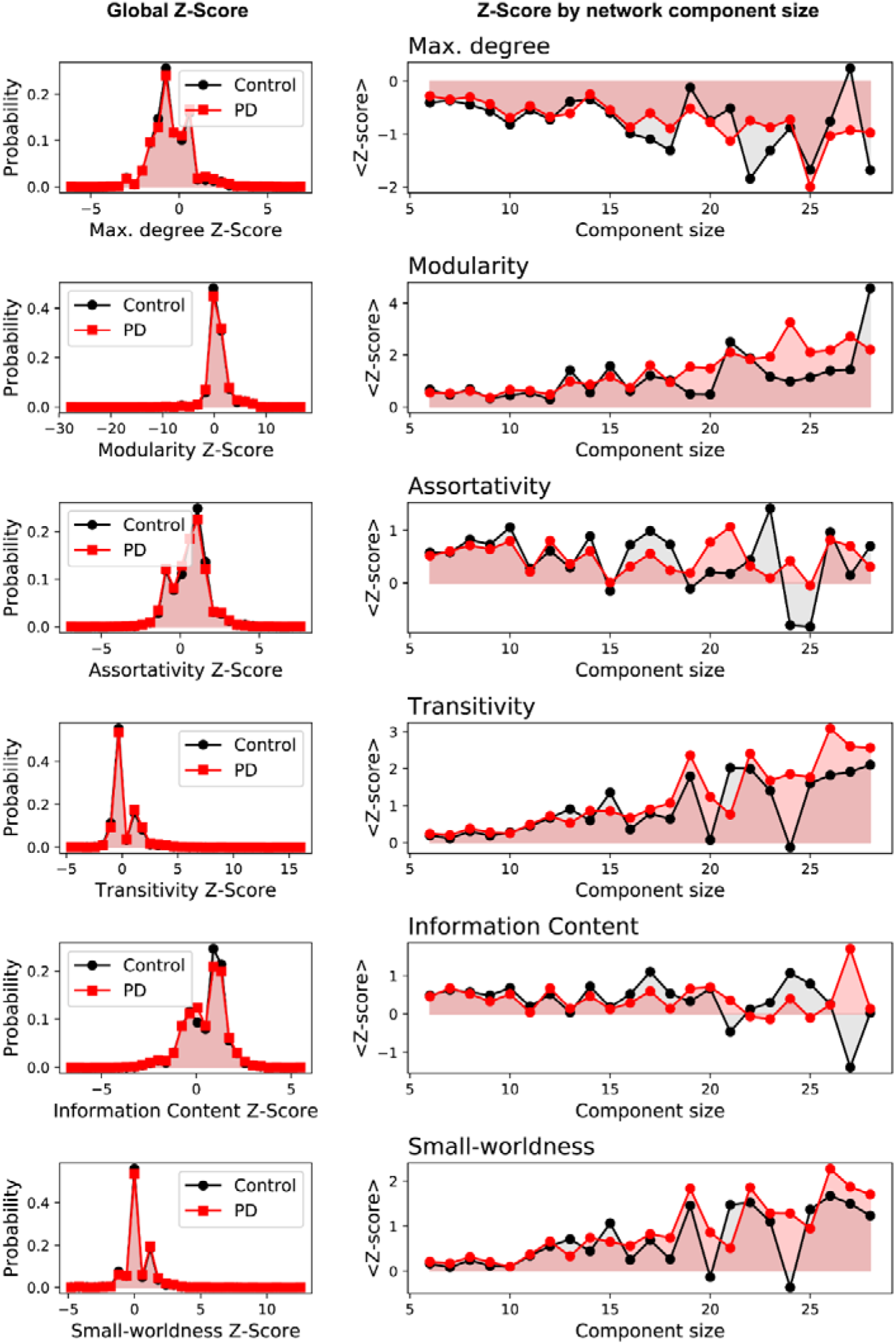
Various graph metrics of mitochondrial interaction networks (MINs) from enteric ganglia of PD. Left, probability of different Z-score of the global MINs from PD patients or healthy controls were plotted; right, Z-score of a given graph metric for the given size of the MIN network components from PD patients or healthy controls. The corresponding statistic results of the graph metrics for the global MINs (refer to Methods) were displayed in **Fig.1f**.

**Supplementary Figure 2.**
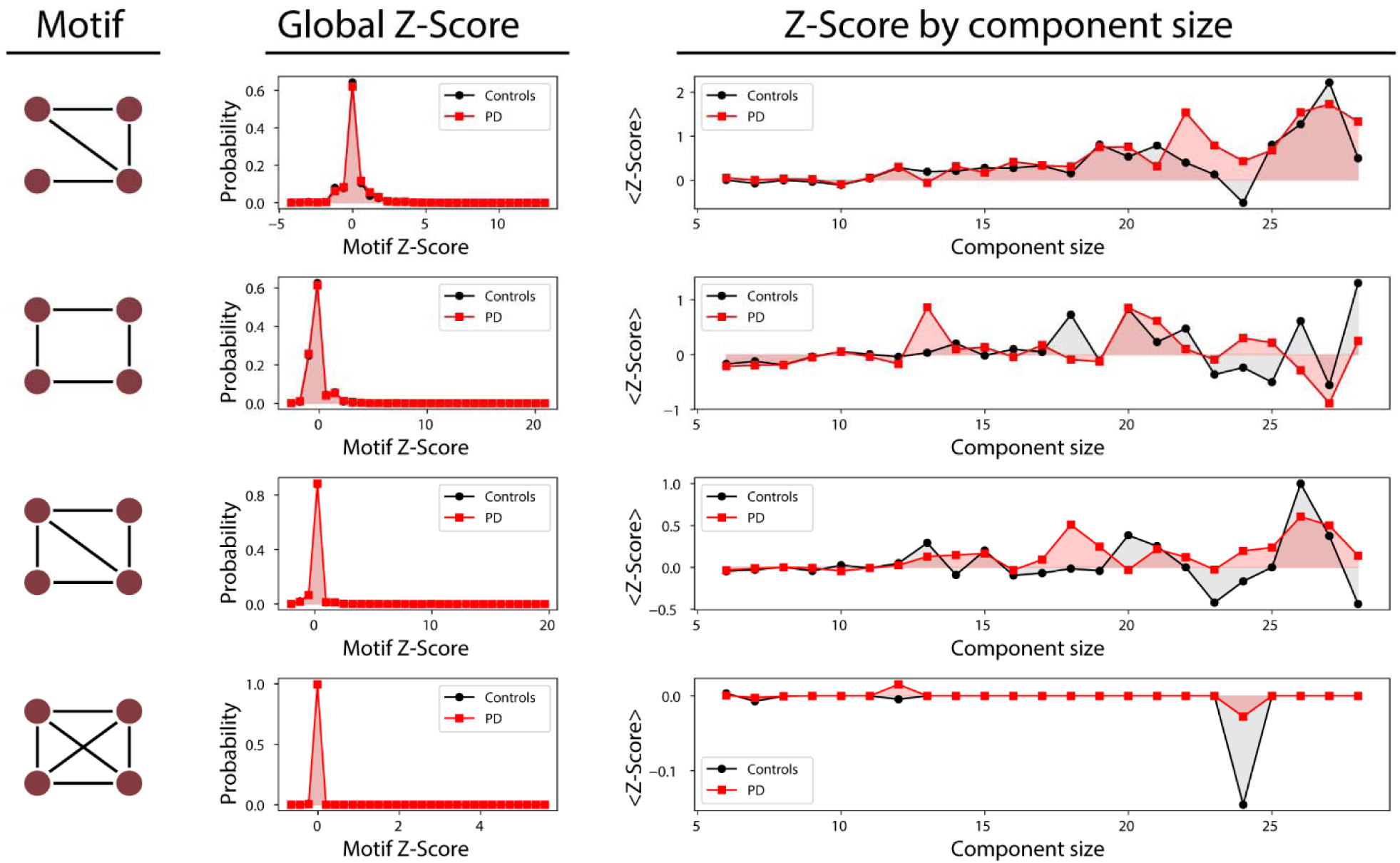
Occurrence of the other types of 4-node motifs within enteric ganglia MINs of PD patients and healthy controls. Each row presents information of the particular type of 4-node motifs from PD or healthy controls. The legend of the display is the same as in **Fig. 2e**.

**Supplementary Figure 3.**
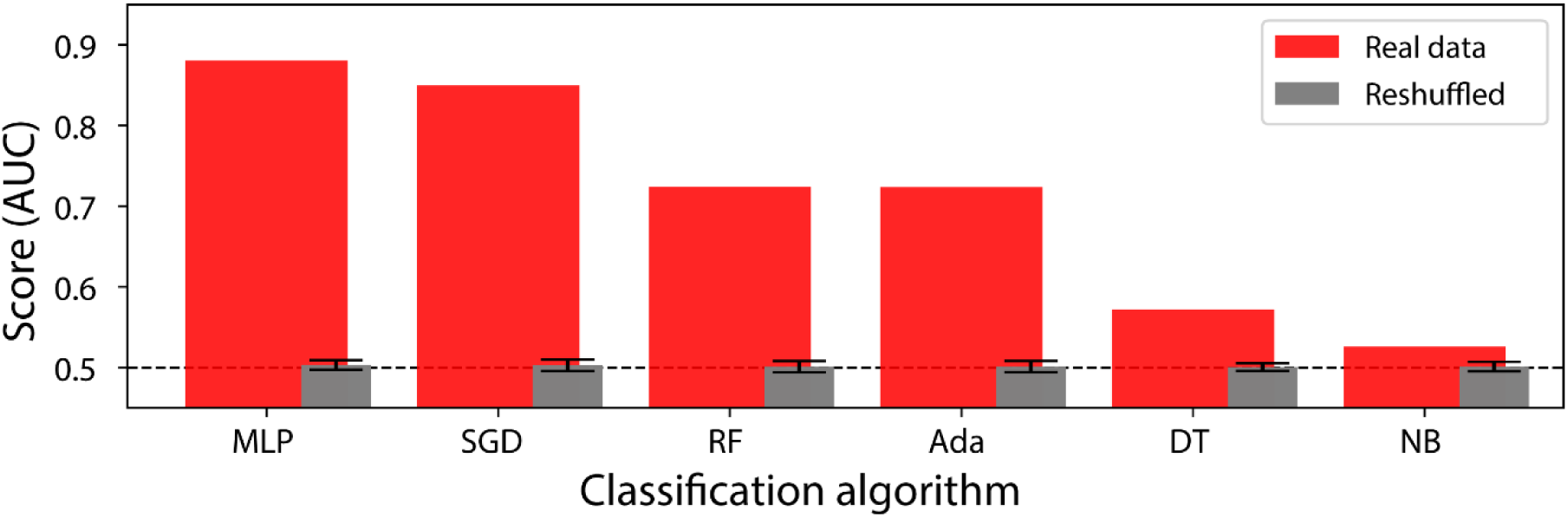
Classification performance of different machine-learning approaches using all the MIN features from all the types of enteric ganglia samples versus randomly-reshuffled datasets. SGD, Stochastic Gradient Descent; MLP, Multilayer Perception; RF, Random Forest; ADA, Ada-boost; DT, Decision Tree; NB, Naive Bayes. The legend ‘Reshuffled’ indicates that we randomly reshuffled the labels 50 times among our samples to test whether we can obtain similar high AUC results in the randomized datasets. The error bar in reshuffled data represents standard deviation.

**Supplementary Figure 4.**
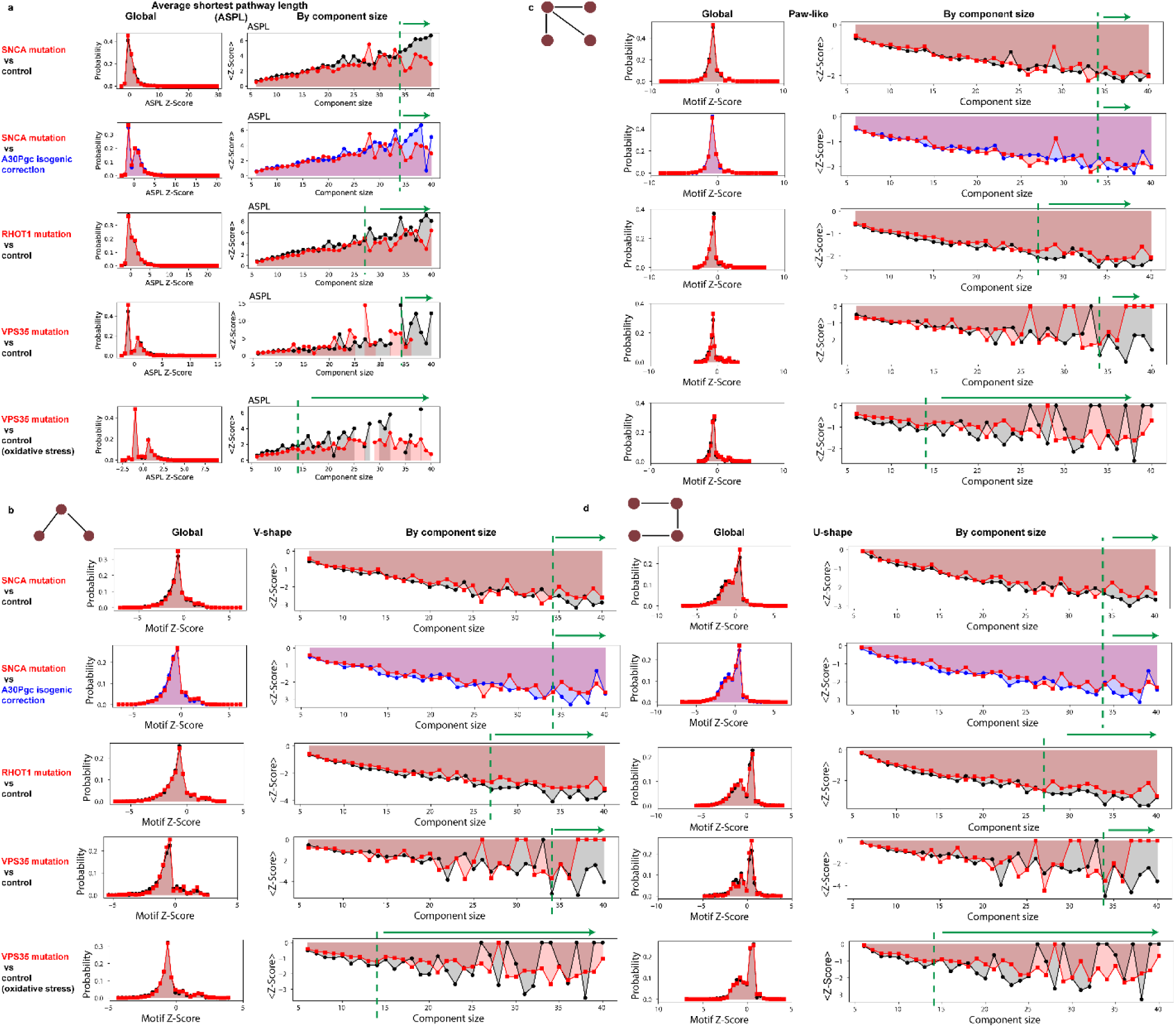
Extended characterization of the MINs of iPSCs-differentiated mDANs of different genetic PD patients. **a**, Distribution of Z-scores of average shortest pathway length (ASPL). Left, histograms representing the probability distribution of the corresponding Z-score from the global MINs of mDANs derived from different patients or matched controls; The right graphs depicting the *Z*-scores for the given size of the MIN components/subnetworks. The green dashed line and the green arrow above the plots highlight the large components that show a clear difference. Of note, mDANs from the VPS35-mutated patient and the matched controls were differentiated with or without (w/o) anti-oxidants (as indicated), while the others were all differentiated in the presence of anti-oxidants. **b, c, d**, Distribution of V-shape motifs (**b**), paw-like 4-node motifs (**c**) and U-shape 4-node motifs (**d**) of the MINs of mDANs derived from different genetic PD patients or matched controls; Left, the histograms representing the probability distribution of the corresponding Z-scores of global MINs from different genetic PD patients or matched controls; the right graphs depicting the *Z*-scores for various sizes of the network components. The green dashed line and the green arrow above the plots highlight the large components that show a clear difference.

## Notes

https://github.com/FengHe001/Mitochondria-network-analysis

https://github.com/FengHe001/Network-matrix-extraction

